# Social isolation reduces oxygen consumption, speed of movement and social preference in wild-type Zebrafish (*Danio rerio*)

**DOI:** 10.1101/2024.11.22.624675

**Authors:** Arpita Ghosh, Aakash Madhav Rao, Prarthna Middha, Atri Bhattacharya, Palakshi Jagetiya, Aryan Tiwari, Shambhavi Rai, Bittu Kaveri Rajaraman

**Affiliations:** Department of Psychology, Ashoka University, Haryana, India; Department of Psychology, University of Alberta, Alberta, Canada; Department of Computer Science, University of Warwick, Coventry, United Kingdom; Department of Computer Science, Ashoka University, Haryana, India; Department of Biology, Ashoka University, Haryana, India

**Keywords:** oxygen consumption, conspecific social preference, average speed, movement

## Abstract

Group living is a common feature of many animals and serves a variety of functions, including predator avoidance, collective foraging, and reproduction. Individuals from group-living species may face isolation from their social group in their natural environment. Prior research has shown that social isolation in zebrafish (*Danio rerio*), a highly social fish, results in physiological and behavioural changes, including reduced locomotor activity and loss of interest in their environment. We investigated whether social isolation was associated with differences in dissolved oxygen consumption rate, movement speed, and social preference in adult zebrafish. We subjected fish to one month of visual and olfactory social isolation, and measured metabolic rate via dissolved oxygen consumption rate before and after social isolation. A control group was not subjected to any form of social isolation but held in comparable housing. We measured movement speed and social interaction before and after social isolation using a social preference assay. We found that socially isolated fish show a significant decrease in oxygen consumption rate compared to Day 0 isolation baseline levels, whereas control fish did not show this decrease. After a month of social isolation fish also showed lower social preference, and lower movement speed. These findings could suggest reduced energy expenditure in the context of social isolation in this species. This work enhances our understanding of the physiological and behavioural consequences of isolation in social animals.

## Introduction

Zebrafish (*Danio rerio*), like humans, are highly social creatures that form social hierarchies and exhibit behaviours such as shoaling and schooling with others of their species (Fontana et al., 2022). Group behaviour is posited to help escape predators and enhance foraging and mating success (Paul et al., 2010; Fontana et al 2022; Mukherjee et al., 2022). When zebrafish are experimentally isolated from their conspecifics, they exhibit consequences such as reduced locomotor activity, reduced interest in their environment, decreased foraging, and changes in serotonergic and dopaminergic pathways associated with mood regulation (Zellner et al., 2011; Marco et al., 2016; Shams et al., 2017; Tunbak et al., 2020; Lachowicz et al., 2021). Social isolation is an ecologically relevant paradigm for zebrafish, which live in rain-fed streams that can dry up to form shrinking ponds, which can later be reconnected by heavy rains (Engeszer et al., 2007; Arunachalam et al., 2013). Long-term social isolation significantly reduces social preference and locomotor activity in zebrafish, whereas brief periods have less pronounced effects on behaviour (Zellner et al., 2011; Shams et al., 2017; Singh et al., 2026).

An essential mechanism through which social isolation and reduced movement can affect survival is their impact on metabolic rate. The stress of social isolation (Daniel and Bhat, 2022) could increase metabolic rates as observed in several species such as subordinate great tits (*Parus major*), deer mice (*Peromyscus maniculatus*), dippers *(Cinclus cinclus*), and Atlantic Salmon (*Salmo salar*) among others (Provost et al., 2022; Fernandes et al., 2023; Zhang et al., 2025), trading off against the possible metabolic savings due to reduced movement demonstrated in carp (*Cyprinus carpio L.*) and roach (*Rutilus rutilus (L.*) (Ohlberger et al., 2005). Metabolic savings have been shown to lead to improved survival by reducing aging through lowering oxidative stress (Bartke et al., 2020) or by being repurposed towards improved immune systems in mice (Bird, 2019) or reproductive systems in certain fish species (Palstra, Crespo, et al., 2010; Palstra, Schnabel, et al., 2010). The direction of downstream effects on reproductive investment change is also unclear – reduced swimming triggers gamete development in rainbow trout (*Oncorhynchus mykiss)* (Palstra, Crespo, et al., 2010) and the European silver eel (*Anguilla anguilla*) (Palstra, Schnabel, et al., 2010). In contrast, reduced mating opportunities lead to sperm ageing and reduced sperm quality in zebrafish (Cattelan & Gasparini, 2021) or reduced sperm quantity in guppies *(Poecilia reticulata)* (Cattelan & Pilastro, 2018).

In the present study, we investigated whether social isolation was associated with changes in dissolved oxygen consumption rate and locomotor activity. In aquatic species, metabolic activity is commonly assessed using respirometry, where faster depletion of dissolved oxygen in closed water tanks indicates higher metabolic rate (Cech & Brauner, 2011; Chabot, McKenzie & Craig, 2016).

Using adult wild-type zebrafish, a powerful model system for investigating the neural basis of behaviour through genetic, developmental, and functional neuroimaging tools (Howe et al., 2013; Mari-Beffa, Mesa-Román & Duran, 2021), we quantified changes in dissolved oxygen consumption rate, social preference, and locomotor activity across a period of social isolation. We hypothesised that zebrafish exposed to social isolation would show lower dissolved oxygen consumption rate, reduced locomotion, and decreased preference for conspecific social interaction.

## Methodology

### Subjects

This quantitative experimental study was conducted using adult wild-type zebrafish aged 6–12 months. All fish were procured from a pet store in Daryaganj, New Delhi, India, sourced from local water bodies. The fish were housed collectively, during acclimation to the lab for two weeks before the experiment, in a ZebTec Active Blue - Stand Alone system (Tecniplast, PA, USA), which maintains water temperature at 28°C, pH at 7.50–8.50, conductivity at 650–700 µS/cm, and a bio-filter based cleaning system, alongside regular automatic changes in water to maintain low nitrate content. Fish were then transferred into glass holding tanks measuring 30 × 20 × 15 cm, with a water column height of 15 cm. Each tank was divided into six equally sized compartments, each approximately 10 × 10 cm, with one fish housed per compartment and six fish housed per tank. The compartments were separated by transparent barriers, which allowed visual exposure to neighbouring fish. These barriers functioned as physical movement barriers with gaps small enough to prevent fish from crossing into adjacent compartments but sufficient to allow water-borne olfactory cues to pass between compartments. Thus, the barriers maintained visual and olfactory exposure to conspecifics. This housing stage therefore provided physical separation without sensory isolation. From the procured population, 60 fish were randomly selected and transferred to the above-mentioned tanks. At the start of the experiment, 30 fish were placed in the social isolation condition and 30 in the control condition. Fish in the social isolation condition were housed individually for 30 days with complete visual and olfactory isolation, whereas control fish remained socially housed in the same housing setup as before for the same duration. Four fish in the social isolation condition did not survive until the end of the isolation period (day 31), leaving 26 socially isolated fish for analysis. Complete paired day 0 and day 31 oxygen consumption data were available for 17 control fish. Therefore, the final oxygen consumption dataset comprised 26 socially isolated fish and 17 control fish. They were fed ad libitum Tetra-Tetramin flakes once daily (Singh et al., 2026). All experiments were conducted between 10:00 a.m. and 4:00 p.m.

A designated set of size and age-matched 20 fish (10 females and 10 males) was communally housed in separate glass tanks of the same dimensions as the experimental holding tanks (30 × 15 × 20 cm), but without internal barriers. Each tank contained water from the same recirculating housing system. Four fish were randomly selected from this group to serve as the shoal in the social preference tests. We maintained these 20 fish throughout all experiments to ensure consistency in the shoal composition.

The care and use of animals followed national and institutional guidelines, including the Guidelines of CPCSEA for Experimentation on Fishes (CPCSEA, 2021). Ethical approval was obtained from Ashoka University’s Institutional Animal Ethics Committee (approval no. ASHOKA/IAEC/2/2022/6). The study was planned with reference to the PREPARE guidelines (Smith et al., 2018) and reported in accordance with the ARRIVE 2.0 guidelines (Percie du Sert et al., 2020). The manuscript was also prepared in accordance with the ASAB/ABS guidelines for the ethical treatment of nonhuman animals in behavioural research and teaching (ASAB Ethical Committee/ABS Animal Care Committee, 2023).

### Experimental Timeline

The study used a mixed repeated-measures design for dissolved oxygen consumption, with phase (day 0 and day 31) as the within-subjects factor and group (social isolation and control) as the between-subjects factor. The same fish were assessed before and after the experimental period, allowing the dissolved oxygen consumption rate to be compared between socially isolated and control fish across the 31 day time period.

On day 0, we first recorded the social preference alongside the movement rate of the fish. Subsequently, following a 15-minute rest period, the dissolved oxygen consumption rate of the experimental fish was recorded. The experimental fish was placed in social isolation immediately following the cessation of the dissolved oxygen consumption rate recording. We ensured that animal transfer occurred promptly and carefully, using sterilised fresh nets to minimise the transfer of conspecific olfactory cues and minimise transfer-induced stress. We then kept the experimental fish in their isolation chamber for 30 days. On the 31st day, we recorded the social preference and dissolved oxygen consumption rate in the same manner as on day 0. Behavioural outcomes, including speed and social preference, were assessed in the social isolation group before and after the isolation period in a within-subjects design. These behavioural outcomes were utilized as a manipulation check, since changes in locomotor activity and conspecific preference in socially isolated zebrafish are already well established. Socially reared zebrafish have been reported to show higher movement activity than socially isolated fish during both acclimation and social viewing periods (Tunbak et al., 2020), and group-reared zebrafish larvae show greater locomotor activity than individually reared larvae (Zellner et al., 2011). Similarly, zebrafish show robust conspecific preference and shoaling behaviour under social housing conditions (Pham et al., 2012; Ogi et al., 2021).

Additionally, we also ran control trials for dissolved oxygen rate measured among socially housed fish (*n* = 17). Control fish were matched as closely as possible to the socially isolated fish in strain, age, body size, housing conditions, and experimental handling. On day 0, dissolved oxygen consumption rate was measured among fish obtained from community tanks, using the same protocol used for socially isolated fish as detailed above. After the day 0 measurement, the subjects were returned back to the community tank, and on day 31, their dissolved oxygen consumption rate was measured again.

### Operationalisation and Experimental Design

#### Social Isolation

Before isolation, fish were physically separated but retained visual and olfactory contact with conspecifics. The sex of the fish was noted (14 males, 12 females). During isolation, each fish was transferred to an individual plastic container (height = 30 cm, diameter = 10 cm, water column height = 15 cm), where exposure to both visual and water-borne olfactory cues from conspecifics was prevented. Each container was covered to prevent visual exposure to conspecifics. To prevent exposure to water-borne conspecific olfactory cues and to standardise the rearing medium, fish were maintained in freshly prepared embryo water throughout the isolation period. Embryo water was prepared from a stock solution following Cold Spring Harbor Laboratory guidelines (Williams & Renquist, 2016) and diluted by adding 16.5 mL of stock solution to 1 L of distilled water for each isolated fish (see Appendix E). The stock solution was stored in a climate-controlled environment at approximately 28°C. To maintain water quality, 50% of the water in each isolation container was replaced twice weekly using freshly prepared embryo water. Fish were isolated for 30 days. This duration was selected as an intermediate isolation period relative to previously reported zebrafish social isolation protocols, which range from 24 hours to 6 months (Shams, Amlani, et al., 2017), while remaining feasible within the logistical constraints of this study (see Appendix D).

#### Dissolved Oxygen Consumption

Our custom experiment set-up (Figure 1A) featured a large external chamber that was uniformly illuminated and connected to a temperature control system. A water inlet and outlet facilitated circulation within the outer chamber, maintaining the desired temperature. Inside this outer chamber was a smaller inner chamber with a capacity of approximately 100 mL, which served as the testing environment for the experimental fish.

**Figure 1:**
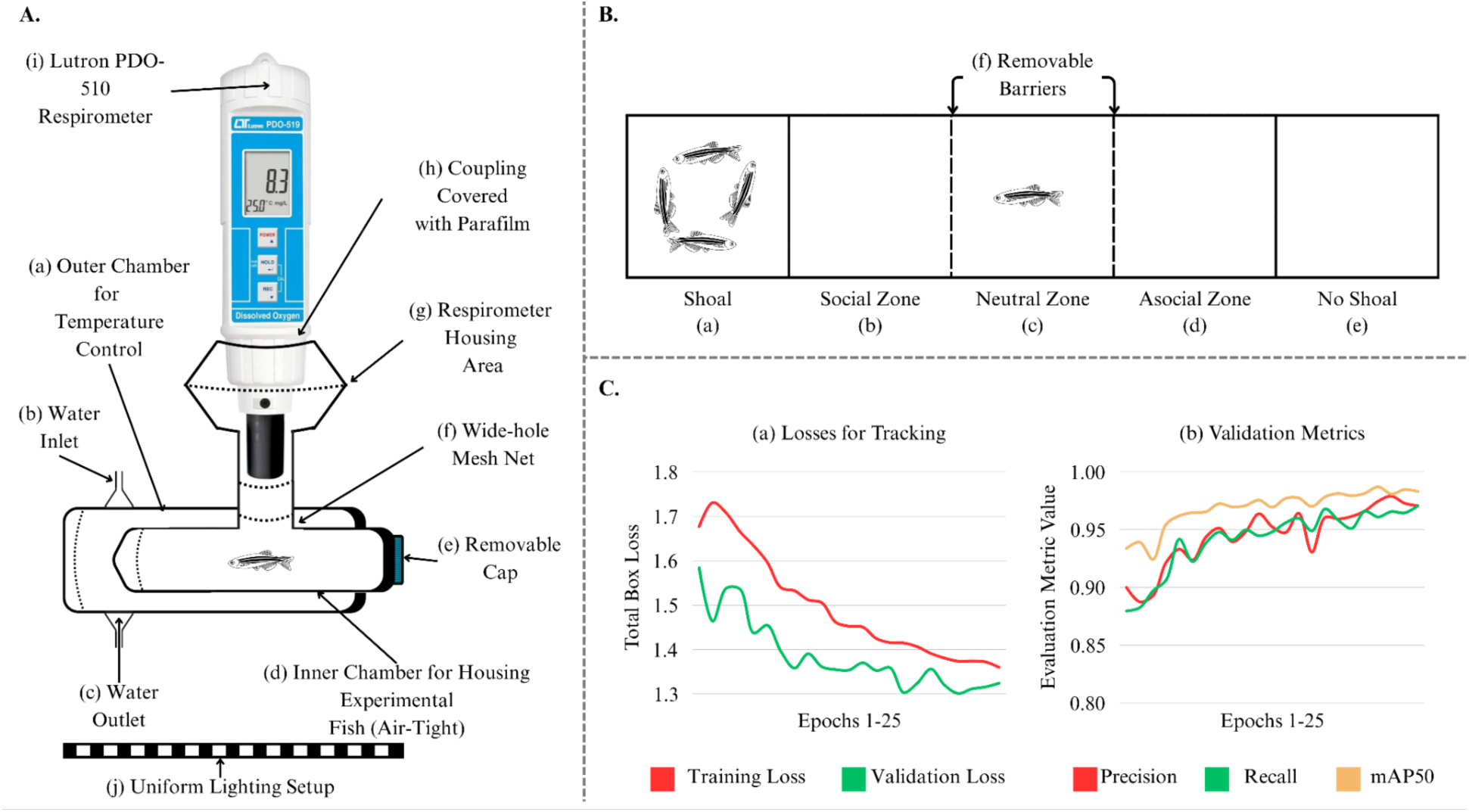
(A) Dissolved oxygen consumption measurement apparatus with uniform lighting, (B) Social preference measurement setup with two shoal positions and three behavioural zones separated by removable barriers, (C) Automated fish movement tracking system losses and metrics.

We introduced an experimental fish into the inner chamber through the upper respirometer housing slot to begin the experiment. Next, we placed a wide-holed net into the same slot to prevent the fish from swimming upward and having direct contact with the probe of our respirometer. Following this, we mounted the respirometer - a Lutron PDO-519 dissolved oxygen meter - into the designated housing, sealing the connection with parafilm to ensure the inner chamber was airtight. This configuration allowed no water exchange between the two chambers, guaranteeing accurate dissolved oxygen measurements. The variation in dissolved oxygen levels over time directly indicated the oxygen consumption of the experimental fish, thereby providing a means to measure its dissolved oxygen consumption rate (Voutilainen, Seppänen & Huuskonen, 2011; Francis-Floyd, 2020; Divakaruni & Jastroch, 2022).

Once we placed the parafilm to seal the inner chamber, we began recording readings as reflected by the respirometer. We checked the reading on the respirometer every minute and noted it down every five minutes for a total of 90 minutes. The 90-minute measurement duration was selected based on pilot trials and fish welfare considerations. During pilot measurements, dissolved oxygen concentrations were consistently maintained above 3.0 mg/L until the 90-minute mark; therefore, 90 minutes was set as a precautionary endpoint to allow a measurable decline in dissolved oxygen while avoiding prolonged exposure to hypoxic conditions. The 90-minute assay was terminated before dissolved oxygen levels fell below 3.0 mg/L. This threshold was selected as a conservative cut-off, as 3.0 ± 0.3 mg/L has been classified as mild hypoxia in zebrafish, while lower levels such as 1.0 ± 0.2 mg/L are associated with severe hypoxic exposure and oxidative damage (Feng et al., 2016). This remains more conservative than Rees, Sudradjat, and Love (2001), who reported mild and severe hypoxia below 0.8 and 0.4 mg/L respectively. In the present study, mean endpoint dissolved oxygen concentrations remained well above this mild-hypoxia threshold across groups: 4.07 ± 0.19 mg/L in Day 0 control, 3.30 ± 0.28 mg/L in post-control, 3.71 ± 0.28 mg/L in Day 0 isolation, and 3.47 ± 0.12 mg/L in post-isolation measurements. Thus, the 90-minute endpoint allowed reliable estimation of oxygen consumption rate while minimizing the risk of exposing fish to more severe oxygen depletion.

To account for the possibility that dissolved oxygen consumption rate may change over time independently of social isolation, we measured dissolved oxygen consumption in a time-matched control group (*n* = 17) of zebrafish maintained under comparable conditions in socially housed tanks. These fish were measured across the same interval as the socially isolated fish. Additionally, as body size can influence dissolved oxygen consumption rate, body weight was also measured in batches of socially isolated (*n* = 20) and control fish (*n* = 43) at Day 0 and Day 31 (Goldson et al., 2026).

#### Social Preference and Movement Speed

We defined social preference as the tendency to live near and interact with conspecifics, as well as the motivation to identify and approach them (Liu et al., 2016). Social preference was measured as the amount of time the experimental fish spent near conspecifics relative to the time spent away from them.

For this assay, we used a uniformly lit 50 × 10 × 12 cm plexiglass tank with a 10 cm water column, divided laterally into five equal chambers (Figure 1B). All four sides of the tank were covered with an opaque black membrane to prevent external confounding stimuli. The tank contained four transparent barriers, two fixed and two removable (Pham et al., 2012). The fixed barriers were not watertight, allowing all fish to share the same water.

A shoal of four adult display zebrafish was placed in one terminal zone, while the opposite terminal zone was left empty, corresponding to zones (a) and (e) in Figure 1B. During the acclimatisation phase, the experimental fish was placed in the neutral zone, marked as (c) in Figure 1B, and allowed to acclimate for 30 s (Pham et al., 2012). During the test phase, the removable barriers were removed, and the experimental fish was allowed to explore the tank for 300 s (Pham et al., 2012). The acclimatisation and test-phase durations were informed by Ogi et al. (2021) and matched the modal durations reported across the 28 studies evaluated in that review. All trials were recorded in MP4 format using a Lenovo 300 FHD 1080P Megapixel webcam. The camera was mounted on a tripod 60 cm above the bottom of the tank to capture a complete top-view recording.

The zone adjacent to the shoal was defined as the social zone, whereas the zone at the opposite end of the tank was defined as the asocial zone; these correspond to zones (b) and (d) in Figure 1B. The central chamber was defined as the neutral zone. For each trial, we quantified the time spent by the experimental fish in the social, neutral, and asocial zones during the 300-s test phase. As the shoal position was alternated between trials to control for lateral bias, zone identity was defined relative to the shoal location in each trial.

#### YOLO-Based Fish Tracking

We converted each experimental fish video into grayscale and cropped it to include only the three central zones (see (b), (c), and (d) in Figure 1B). Subsequently, the videos were stored as .mp4 files and cropped to a fixed duration of 5 minutes, starting from the moment the removable barriers were raised, eliminating any extra recording time.

To train YOLOv8 (Varghese & Sambath, 2024), approximately 50 frames were sampled from each recorded behavioural test video, yielding approximately 6,000 frames. Frames were annotated using Roboflow (Dwyer & Nelson, 2022) and split into training, test, and validation datasets using a 70:20:10 ratio (Tan et al., 2021). The model was trained to detect and localise the experimental fish using bounding boxes (Redmon et al., 2016).

Object detection was performed using the pretrained YOLOv8m model. Training was conducted for 23 epochs, with a batch size of 8 and an input image size of 640 × 640 pixels. The model was trained using pretrained weights, automatic optimiser selection, an initial learning rate of 0.01, final learning-rate factor of 0.01, momentum of 0.937, and weight decay of 0.0005. A warm-up period of 3 epochs was used. Training was performed on Apple Silicon using the MPS device, with automatic mixed precision enabled.

As the model was trained for single-object bounding-box localisation rather than image-level classification, model performance was evaluated using object-detection metrics rather than percentage accuracy. Specifically, we report precision, recall, and mean average precision at an intersection-over-union threshold of 0.5, hereafter mAP50 (Henderson & Ferrari, 2016). Training and validation loss curves, along with precision, recall, and mAP50 across epochs, are shown in Figure 1C. The model achieved an mAP50 of 0.985, indicating strong detection performance for identifying and localising the experimental fish. The trained model was then used to identify the position of the experimental fish in each frame, and the resulting positional coordinates were used for downstream analyses of zone occupancy and movement.

## Statistical Analysis

### Dissolved Oxygen Consumption

We recorded oxygen consumption rate every five minutes for 90 minutes in experimental fish (*n* = 26) and control fish (*n* = 17). Experimental fish were tested on day 0, socially isolated for 30 days, and tested again on day 31. Control fish were tested on the same schedule but remained socially housed throughout the 30-day period. The experimental and control groups, therefore, contributed 18 time-points of measurement each at the 0th and 31st day. Dissolved oxygen consumption rate was estimated separately for each fish and day by fitting a linear slope across the 18 dissolved oxygen measurements. Slopes were multiplied by −1 so that higher values indicated faster dissolved oxygen consumption. For each fish, a change score was calculated as the day 31 oxygen consumption rate minus the day 0 oxygen consumption rate. In experimental fish, this represented change following 30 days of social isolation; in control fish, it represented change across the same 30-day period while fish remained socially housed. Normality of these change scores was assessed using Shapiro–Wilk tests, and equality of variance between groups was assessed using Levene’s test. Wherever the assumption of equal variances was not met, Welch’s independent-samples t-test was used as the primary analysis to compare the change in oxygen consumption rate between groups. Mann–Whitney U, permutation testing, and bootstrap confidence intervals were used as sensitivity analyses. All statistical analyses were conducted in Python.

### Social Interaction and Movement Speed

Zone occupancy and movement speed were quantified using the positional information obtained during the social preference test. For each trial, the total time spent by each fish in the social, neutral, and asocial zones was calculated and expressed as a percentage of the total test duration. Since each fish completed two side-swapped trials, the proportion of time spent in each functional zone was averaged across trials to obtain one fish-level estimate for each zone.

As time spent in the three zones was constrained to sum to 100%, social and asocial zone times were not treated as independent outcomes. Instead, social preference was quantified using a social preference index (SPI), calculated as:

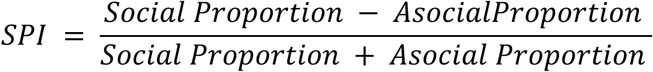

Higher SPI values indicate stronger preference for the shoal zone. A Day 0 and Day 31 SPI value was calculated for each fish, and the change in SPI was calculated as Day 31 minus Day 0.

Movement speed was calculated from the same positional data. For each trial, the total distance covered was estimated by summing the Euclidean distance between successive x-y coordinates across the full test phase. Average speed was then calculated by dividing the total distance by the trial duration. As each fish completed two side-swapped trials, distance was summed across trials and divided by the total duration of both trials to obtain one fish-level estimate of average movement speed before and after social isolation. Paired change in movement speed was calculated as Day 31 average speed minus Day 0 average speed.

Normality of paired change scores was assessed visually using QQ plots (see Appendix C). For non-normally distributed data such as SPI, changes before (Day 0) versus after (Day 31) social isolation were analysed using a Wilcoxon signed-rank test. A paired-samples t-test was also reported as a sensitivity analysis. For normally distributed data, such as movement speed changes in average speed before and after isolation, were analysed using a paired-samples t-test. A Wilcoxon signed-rank test and a paired sign-flipping permutation test were reported as sensitivity analyses. Zone-specific changes in social, neutral, and asocial occupancy were reported descriptively as percentage-point changes, because the three zone proportions are not statistically independent.

## Results

### Dissolved Oxygen Consumption Rate

Using the computed slope-derived estimates, raw dissolved oxygen trajectories showed different slopes of decline between control and socially isolated fish across days of testing (Figure 2A). For visualisation, baseline-centred trajectories were computed by subtracting each fish’s mean Day 0 dissolved oxygen value from all of its Day 0 and Day 31 dissolved oxygen measurements. These baseline-centred trajectories showed the same pattern as the raw data (Figure 2C), with Day 31 values in the social isolation group remaining higher relative to baseline than their Day 0 trajectory, suggesting a shallower dissolved oxygen decline after isolation.

**Figure 2:**
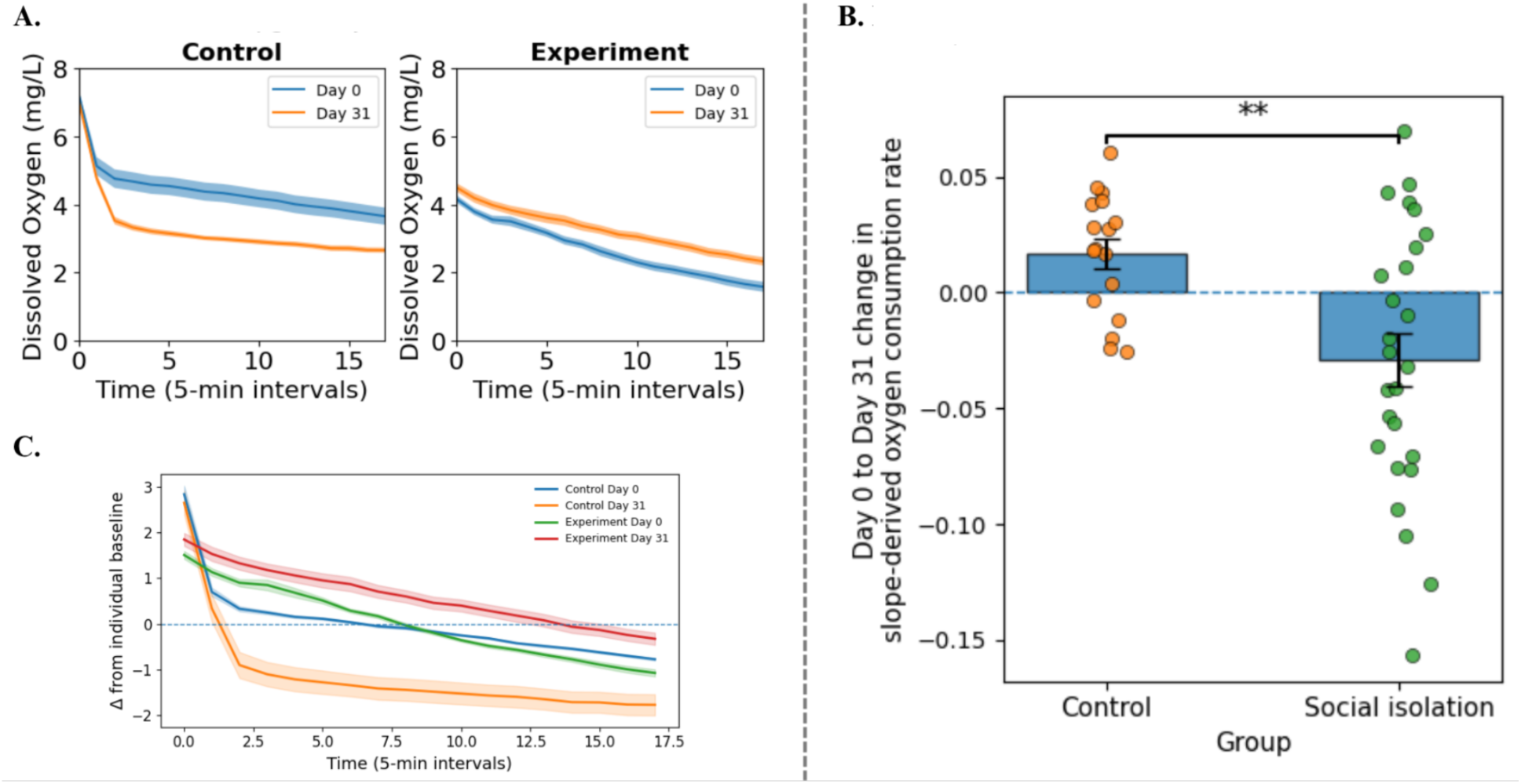
(A) Raw dissolved oxygen trajectories across the groups and intervention (Day 0 and Day 31) and stages, with shaded regions indicating ± SEM. (B) Fish-level Day 0 to Day 31 change in dissolved oxygen consumption rate in control and socially isolated fish. Each point represents one fish, and black markers indicate group means ± SEM. Negative values indicate a reduction in dissolved oxygen consumption rate after social isolation. (C) Baseline centered dissolved oxygen trajectories across the groups and intervention stages (Day 0 and Day 31), with shaded regions indicating ± SEM. (*p < .05, **p < .01, ***p < .001).

**Figure 3:**
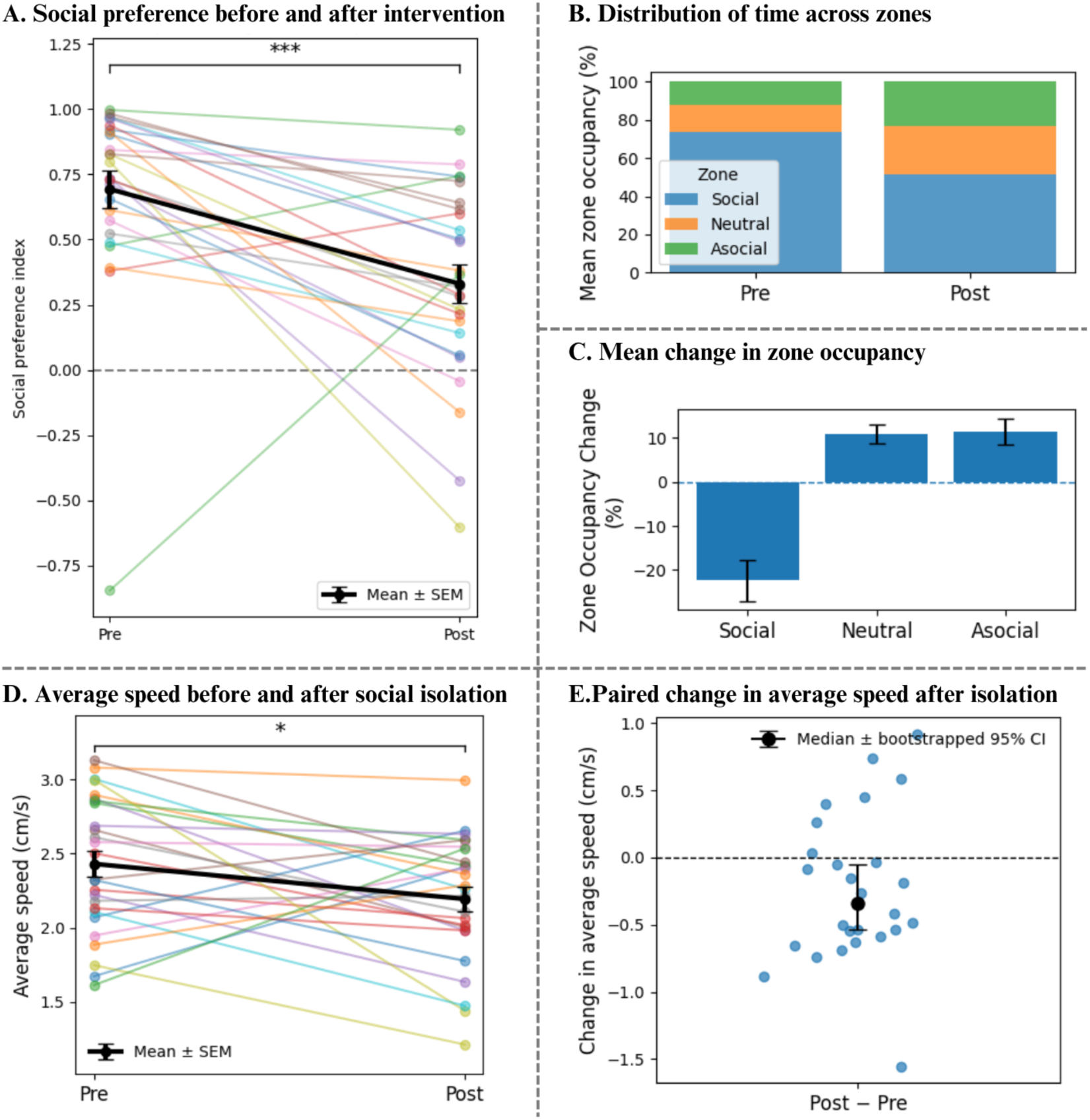
(A) Fish-level social preference index (SPI) before and after social isolation (Day 0 and Day 31). Each coloured line represents one fish, and black points show the group median with error bars indicating the interquartile range. SPI significantly decreased (*p* < 0.001) from Day 0 to Day 31 post-isolation, indicating reduced social preference. (B) Mean percentage of time spent in the social, neutral, and asocial zones before and after social isolation. (C) Mean Day 0 to Day 31 change in zone occupancy for each tank zone. Error bars indicate SEM. Changes are shown in percentage points. Zone-level changes are presented descriptively because the three zone proportions were constrained to sum to 100%. (D) Fish-level average movement speed before and after social isolation. Each coloured line represents one fish, and black points show the group mean with error bars indicating SEM. Average movement speed significantly decreased (*p* = 0.041) from Day 0 to Day 31 post-isolation. (E) Paired Day 0 to Day 31 change in average movement speed for each fish, with random jitter along the x axis to improve visibility. The dashed horizontal line indicates no change. The black point shows the median paired change, and error bars indicate the bootstrapped 95% confidence interval. (*p < .05, **p < .01, ***p < .001).

Fish-level change scores were then calculated as post-isolation minus Day 0 oxygen consumption rate. Control fish showed a small increase in dissolved oxygen consumption rate from Day 0 to Day 31, whereas socially isolated fish showed a decrease (Figure 2B). The mean change was 0.017 (± 0.026) in the control group and −0.029 (± 0.058) in the social isolation group, giving a mean difference in change of −0.046 (± 0.032). Shapiro–Wilk tests did not indicate strong evidence against normality in either group (control: Shapiro-Wilk *W* = 0.944, *p* = 0.3688; Socially Isolated: Shapiro-Wilk *W* = 0.979, *p* = 0.8587); however, Levene’s test showed unequal variances between groups (Levene test: *W* = 10.154, *p* = 0.0028). Therefore, Welch’s independent-samples t-test was used as the primary analysis. This showed that the Day 0 to Day 31 change in dissolved oxygen consumption rate differed significantly between groups, *t* = −3.531, *p* = 0.001. The effect size was large, Cohen’s *d* = 0.956, indicating that socially isolated fish showed a greater reduction in dissolved oxygen consumption rate relative to controls.

Sensitivity analyses supported this result. A Mann–Whitney U test also showed a significant difference between groups, *U* = 112.000, *p* = 0.007. A fish-level permutation test produced a similar result, with an observed mean difference of −0.046 and *p* = 0.003. The bootstrap 95% confidence interval for the mean difference ranged from −0.072 to −0.021, which did not include zero (see Appendix A).

Additionally, to assess whether these differences in oxygen consumption rate could be attributed to differences in body weight, body weight was compared between the Day 0 and Day 31 samples. As individual fish could not be matched across time points, these comparisons were treated as unmatched sample-level analyses rather than within-individual change analyses. Body weight, measured in grams, did not differ detectably between Day 0 and Day 31 in the socially isolated group, despite deviations from normality at both time points (Day 0: *M* = 0.22 g, *SD* = 0.13, Shapiro–Wilk *W* = 0.671, *p* < 0.001; Day 31: *M* = 0.18 g, *SD* = 0.09, Shapiro–Wilk *W* = 0.745, *p* < 0.001; Mann–Whitney *U* = 244.5, *p* = 0.233). This was supported by an independent-samples bootstrap, which estimated a mean difference of −0.0339 g between the Day 31 and Day 0 samples, with a 95% confidence interval spanning zero [−0.1067, 0.0304]. Similarly, body weight did not differ detectably between Day 0 and Day 31 in the control group as well (Day 0: *M* = 0.20 g, *SD* = 0.07, Shapiro–Wilk *W* = 0.943, *p* = 0.007; Day 31: *M* = 0.21 g, *SD* = 0.06, Shapiro–Wilk W = 0.948, p = 0.051; Mann–Whitney *U* = 1173.0, p = 0.362). The independent-samples bootstrap estimated a mean difference of 0.010 g between the Day 31 and Day 0 control samples, with a 95% confidence interval spanning zero [−0.015, 0.036]. These results suggest that group-level differences in oxygen consumption rate were unlikely to be explained by differences in body weight between Day 0 and Day 31 samples.

We also examined sex-based changes in dissolved oxygen consumption change for the socially isolated group. Change scores were approximately normally distributed in both males and females in the socially isolated group, as assessed using Shapiro–Wilk tests (male: *W* = 0.904, p = 0.177; female: *W* = 0.950, *p* = 0.559). Homogeneity of variance was also met, Levene’s test, *F*(1, 24) = 0.024, *p* = 0.878. Change scores did not differ significantly between males and females (males showed a mean change of −0.007 ± 0.052, whereas females showed a mean change of −0.048 ± 0.057, Welch’s *t*(23.88) = 1.90, *p* = 0.070).

### Social Preference and Movement Speed

Social preference was quantified using a social preference index (SPI), calculated as (social zone time − asocial zone time)/(social zone time + asocial zone time), with higher values indicating stronger preference for the shoal zone. SPI decreased from Day 0 to Day 31 post-isolation, indicating reduced social preference following social isolation. The median change in SPI was −0.384, with a bootstrapped 95% confidence interval of [−0.534, −0.207]. A Wilcoxon signed-rank test showed that this decrease was statistically significant, *W* = 41.0, *p* < 0.001. A paired-samples t-test used as a sensitivity check led to the same conclusion, with SPI significantly lower after isolation than before isolation, *t*(25) = 3.72, *p* = 0.001.

Zone-level occupancy patterns supported this reduction in social preference. Fish spent less time in the social zone after social isolation, with a mean decrease of 22.36 percentage points and a median decrease of 27.33 percentage points. This decrease was accompanied by increased occupancy of the neutral and asocial zones. Neutral-zone occupancy increased by a mean of 10.94 percentage points and a median of 13.41 percentage points, while asocial-zone occupancy increased by a mean of 11.41 percentage points and a median of 14.47 percentage points. Since the three zone proportions are definitionally constrained to sum to 100%, these zone-level changes were interpreted descriptively as a redistribution of time across the tank zones rather than as independent outcomes.

Average movement speed was calculated for each fish by summing the distance covered across the two side-swapped trials and dividing this by the total trial duration, with higher values indicating greater movement. Average speed of movement decreased from Day 0 to Day 31 post-isolation, indicating reduced movement following social isolation. Mean speed decreased from 2.43 to 2.19 cm/s, corresponding to a mean paired change of −0.236 cm/s. A paired-samples t-test showed that this decrease was statistically significant, *t*(25) = −2.16, *p* = 0.041, with a small-to-moderate paired effect size, Cohen’s *d* = −0.42. Sensitivity analyses supported the same conclusion: a Wilcoxon signed-rank test was significant, *W* = 94.0, *p* = 0.038, and a paired sign-flipping permutation test also indicated significance, *p* = 0.040. The median paired change was −0.341 cm/s, with a bootstrapped 95% confidence interval of [−0.539, −0.056] (see Appendix D).

## Discussion

Our study examined oxygen consumption rate and social preference in wild-type zebrafish to investigate whether social isolation-induced reductions in movement may be accompanied by bioenergetic changes. By quantifying oxygen consumption alongside changes in movement rate and social preference, this work extends existing literature that has largely focused on behavioural outcomes of social isolation. Following one month of social isolation, zebrafish showed reduced oxygen consumption rate, decreased social preference, and lower movement rates. These findings suggest that social isolation is associated not only with altered social and locomotor behaviour, but also with measurable changes in energetic demand. This approach provides insight into the bioenergetic state of organisms exposed to prolonged social isolation, an aspect that remains relatively understudied compared with behavioural measures alone.

The behavioural changes observed in the present study are consistent with previous work showing that zebrafish exhibit marked behavioural alterations following both acute and chronic social isolation (Lachowicz et al., 2021; de Abreu et al., 2022). Social isolation in zebrafish has been associated with altered locomotor and exploratory behaviour, reduced shoal cohesion, and diminished social preference or social responsiveness (Zellner et al., 2011; Shams et al., 2015, 2017, 2018; Tunbak et al., 2020; Wee et al., 2022). The duration and developmental timing of isolation appear to be important determinants of these effects. Zebrafish reared in complete isolation from birth show significantly lower social preference in adulthood than socially reared individuals (Tunbak et al., 2020), suggesting that prolonged social deprivation can produce persistent reductions in social motivation. Similarly, adult zebrafish acutely isolated for 24 h do not show marked changes in locomotor activity or social preference when re-exposed to conspecifics, whereas individuals isolated for six months exhibit reduced locomotor activity and decreased social preference (Shams, Amlani, et al., 2017). Our findings align with this literature by showing that one month of visual and olfactory social isolation was sufficient to reduce both movement rate and social preference. Importantly, by measuring oxygen consumption alongside these behavioural outcomes, the present study extends previous work by linking isolation-induced behavioural outcomes to a measurable reduction in energetic demand.

As social interaction and locomotion are energetically demanding processes (Ohlberger et al., 2005), the concurrent reduction in movement, social preference, and oxygen consumption rate observed in socially isolated fish suggests a potential decrease in overall energy expenditure. Previous studies have examined these variables largely in isolation, but their findings are consistent with this interpretation. For example, variation in activity levels across fish species, including bonefish, cunner, carp, and roach, has been associated with differences in energy expenditure (Brownscombe, Cooke, & Danylchuk, 2017). Similarly, swimming is energetically more costly than remaining inactive (Ohlberger et al., 2005; Speers-Roesch, Norin & Driedzic, 2018), supporting the broader conclusion that reduced locomotor activity can lower energetic demand (Ohlberger et al., 2005; Millidine, Armstrong & Metcalfe, 2009; Lachowicz et al., 2021). Together, these findings suggest that social isolation-induced reductions in movement and social interaction may be accompanied by a bioenergetically conservative state.

In resource-limited contexts, reduced locomotion may reflect an energy-conserving response. In zebrafish, long-term food deprivation has been shown to produce a chronic decrease in physical activity relative to fed controls, supporting an association between negative energy balance and reduced activity (Novak et al., 2005). Moreover, a high caloric diet in zebrafish has been linked with elevated levels of continuous locomotor activity (Kopp, Legler & Legradi, 2016). Evidence from other teleosts further suggests that the energetic state can shape social behaviour. In juvenile qingbo carp (*Spinibarbus sinensis*), prolonged food deprivation altered sociability, with food-deprived individuals maintaining proximity to shoals, while well-fed individuals with higher standard metabolic rates were less sociable, possibly to reduce competition while meeting higher energetic demands (Killen et al., 2016). Although not directly tested in the present study, similar trade-offs may be relevant in the context of social isolation, where reduced locomotion could constrain foraging activity and thereby limit energy intake under food-limited conditions. These findings, in light of evidence presented in the present study, suggest that reductions in social engagement may, in some contexts, reflect a shift in behavioural priorities under energetic constraints.

Reduced social interaction may also carry potential survival-related benefits in high-risk ecological contexts, particularly under elevated predation pressure. For example, socially withdrawn crustaceans with reduced activity levels have been shown to encounter fewer predators than individuals swimming in shoals, thereby lowering their likelihood of predation by sticklebacks (Ioannou & Krause, 2007). This suggests that reduced activity and social withdrawal may be advantageous in contexts where predator avoidance is prioritised over the benefits of group association. However, these interpretations remain speculative in the context of socially isolated zebrafish, particularly given interspecific differences in ecology, social organisation, and predator-prey dynamics. The present study did not directly measure food intake, energy storage, predation vulnerability or survival. Future investigations should therefore assess whether visual and olfactory social isolation in zebrafish is accompanied by measurable changes in feeding behaviour, body condition, immune activity, motivational state, predation risk, reproductive output, or other fitness-related outcomes.

We must address a few limitations of our study. The small space of the inner chamber in the dissolved oxygen consumption measurement setup could have acted as a stressor for the fish (Rey et al., 2015; Demin et al., 2020). Further because the metabolic assay was conducted in a sealed chamber, declining dissolved oxygen over the 90-minute recording period may have contributed to mild physiological stress, although the assay duration was selected to prevent oxygen levels from falling below acceptable welfare limits (Feng et al., 2016). The central tube had a diameter of 3.7 cm, which is adequately larger than the average adult zebrafish length of 2.8 cm, allowing unrestricted turning and natural horizontal swimming. The dorsal fin remained fully submerged throughout, minimising physical discomfort. However, we acknowledge that the confined space may have posed a mild stressor. While we observed no overt signs of distress, we did not explicitly measure physiological stress markers such as cortisol levels, representing an avenue for future research..

Another potential limitation is that the experimental fish may have briefly seen the experimenter’s hands when removing the removable barriers from the social interaction setup. Although we took care to standardise this procedure across trials using consistent hand movements and timing, visual exposure to the experimenter and sudden water movement may have introduced a mild, uncontrolled stimulus (Clark, Boczek & Ekker, 2011). While the brief exposure was unlikely to have significantly influenced behaviour, future protocols could incorporate visual shielding or automated barrier removal to eliminate this variable. We also anticipated some stress reduction due to the fish experiencing some relief from the visual and olfactory isolation (Faustino, Tacão-Monteiro & Oliveira, 2017; Daniel & Bhat, 2022). Although the metabolic assay itself may impose some handling-related stress, this effect was controlled for by including socially housed fish that underwent the same dissolved oxygen consumption measurements. By comparing Day 0 to Day 31 changes in socially isolated fish with those in socially housed controls, we were able to distinguish changes associated with isolation from changes attributable to repeated testing, handling, or time. Together, these findings suggest that social isolation may shift zebrafish toward a bioenergetically conservative behavioural state, characterised by reduced energetic demand and altered social behaviour. Although the adaptive significance of these changes is likely context dependent, reduced energetic demand may be advantageous under ecological conditions involving resource limitation, pathogen exposure, or predation risk. Future studies should test these physiological and behavioural responses in ecologically relevant contexts to better understand the adaptive significance of social isolation-induced phenotypes.

## Conclusion

In conclusion, this study quantified dissolved oxygen consumption rate, social preference, and movement speed to examine the bioenergetic correlates of behavioural change following social isolation in adult wild-type zebrafish. Across one month of visual and olfactory isolation, isolated fish showed reduced dissolved oxygen consumption rate, lower movement speed, and decreased social preference. The decreased social preference and movement speed are in line with the literature on the effect of 6 months of social isolation, suggesting that a single month of isolation is sufficient to produce these behavioural changes. In addition, we were able to quantify energetic savings in the context of social isolation.

## Author contributions statement

AG and BKR designed the study; AG and AMR made the apparatus; AG, AMR, PM, and SR collected metabolic and movement data; PJ and AT collected weight data; AG and AMR performed data analysis; and AG, BKR, AMR, PM, SR and AT wrote the manuscript.

## Acknowledgements

We would also like to thank Anoushka Damani, Aditi Gudi, Asmi Aggarwal, Ishani Khan, Ishrat Singh, Kashish Bhargava, Keona Gupta, Mahi Pareek, Manya Kocchar, Prarthana Mahajan, Radhika Rajesh Chandwadkar, Reva Sawant, Senna Singh, Sven Aranha, Tanisha Viegas, and Tvisha Tyagi from the Department of Psychology, Ashoka University for their assistance during data collection at various stages.

## Additional Information

We followed CCSEA national and institutional guidelines for the care and use of animals. Ethical approval was obtained from Ashoka University’s Institutional Animal Ethics Committee (approval no. ASHOKA/IAEC/2/2022/6).

## Data Availability

Data and code are available at our project GitHub Repository found at https://github.com/mraoaakash/EcoAdvDep

## Appendices

## Appendix A: Dissolved Oxygen Consumption Rate Measures

**Figure A.1.**
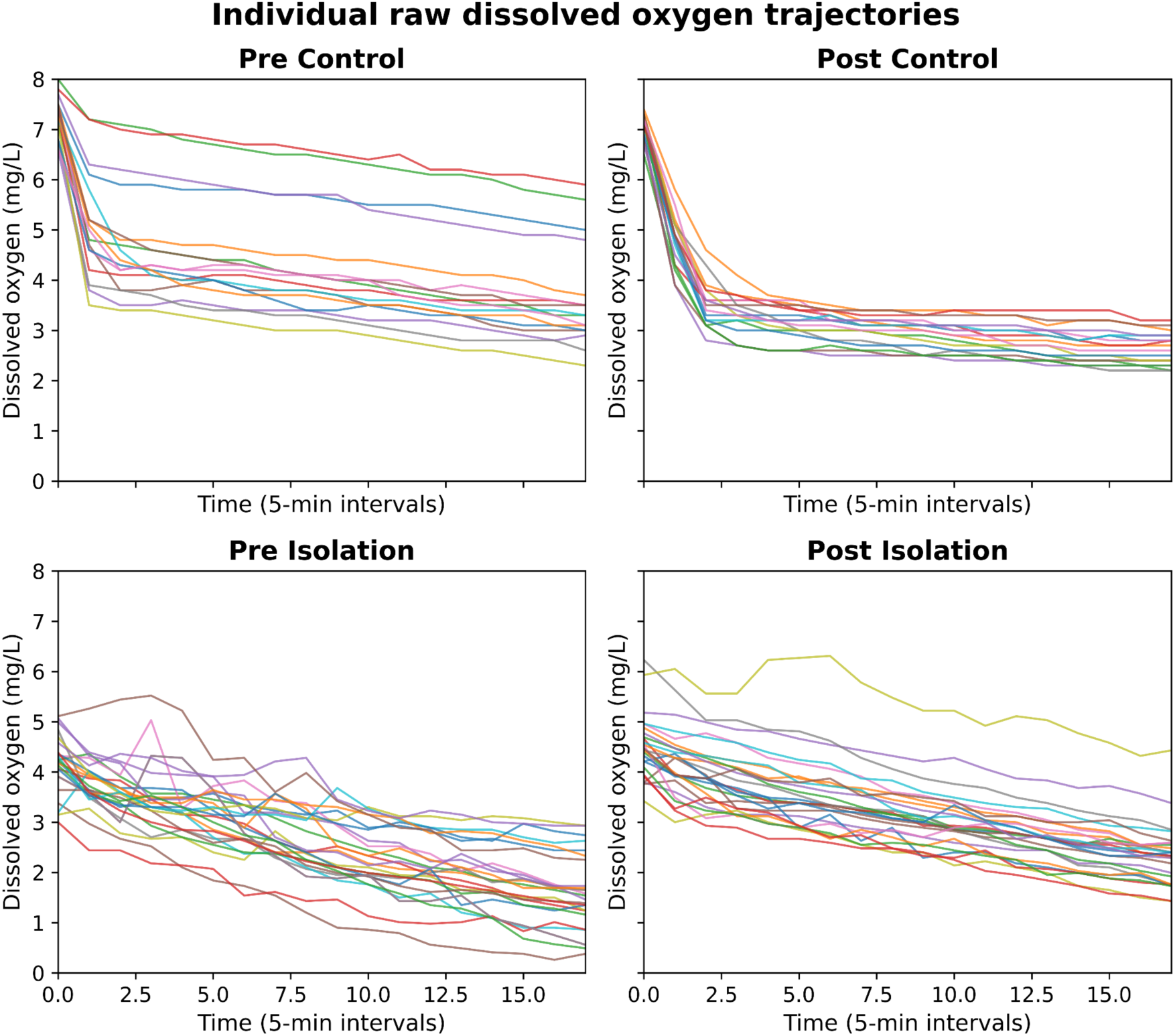
Individual dissolved oxygen trajectories across 18 measurement time-points for Day 0 control, Day 31 control, Day 0 isolation, and Day 31 isolation conditions. Each line represents one fish. This figure shows fish-level variability in dissolved oxygen decline before and after the social isolation.

**Figure A.2.**
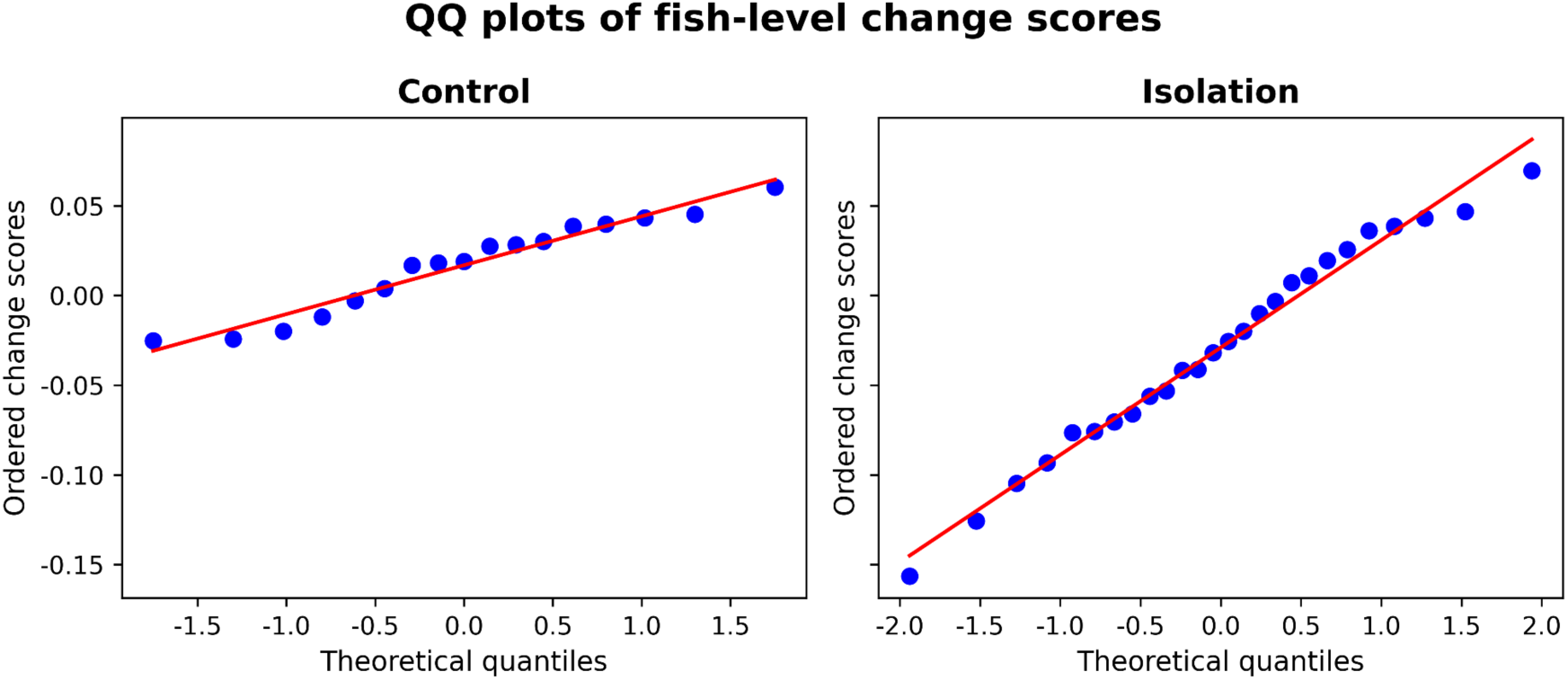
QQ plots showing the distribution of fish-level Day 0 to Day 31 change scores in the control and social isolation groups. These plots were used to visually assess the normality assumption for the change-score analysis.

**Figure A.3.**
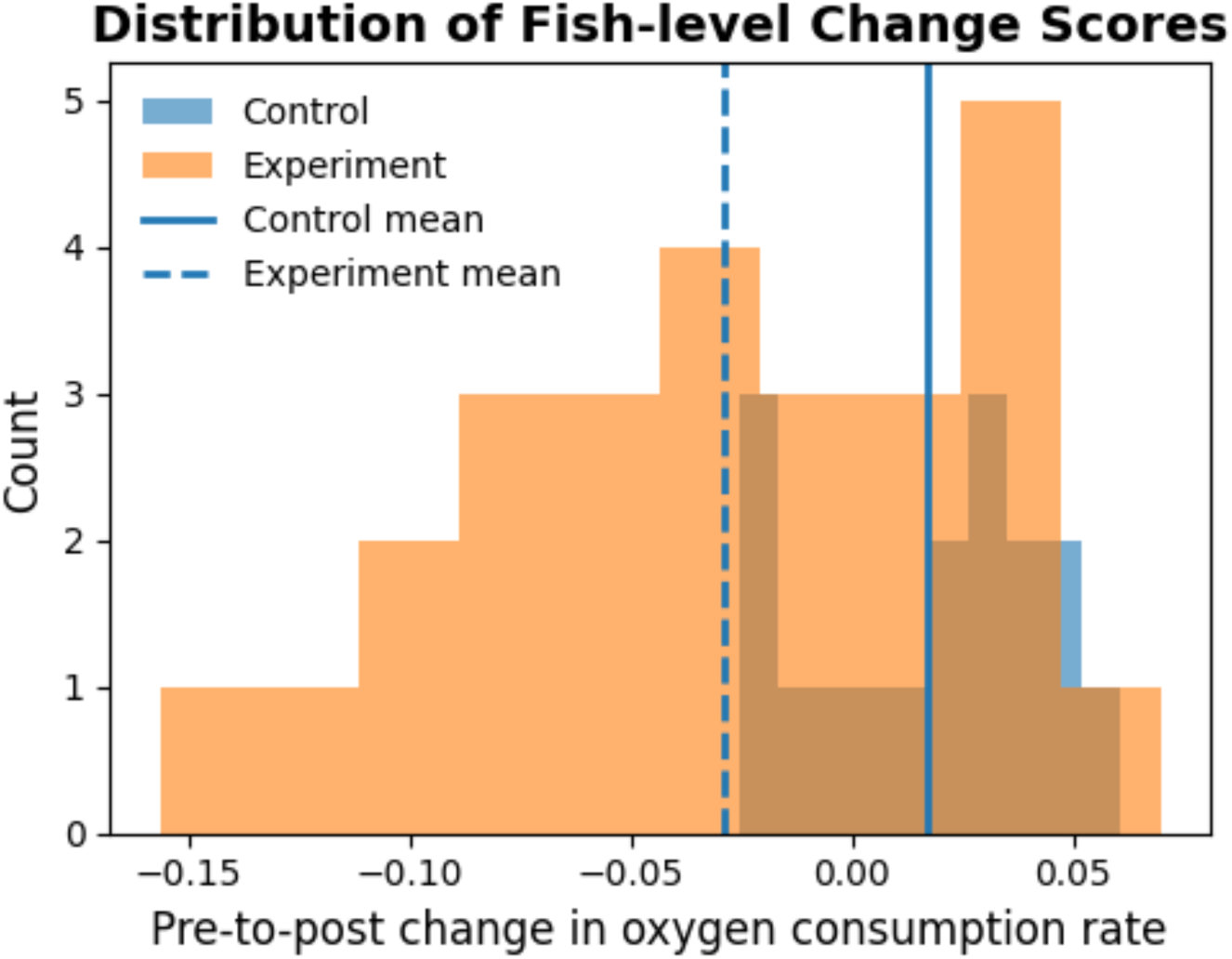
Histogram of Day 0 to Day 31 change in slope-derived dissolved oxygen consumption rate for control and socially isolated fish. Dashed vertical lines indicate group means. Negative values indicate reduced dissolved oxygen consumption rate after social isolation.

## Appendix B: Body-weight Measures

### Socially Isolated Fish

**Figure B.1.**
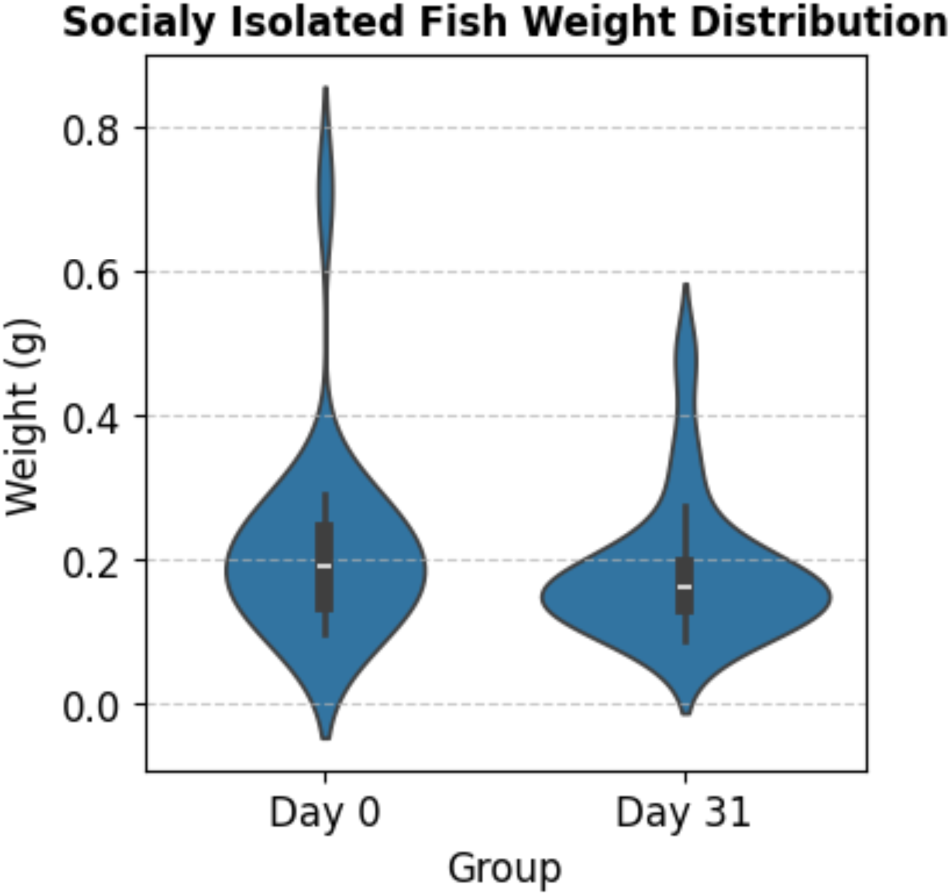
Violin plot for the distribution of body weight in day 0 and day 31 in the socially isolated group.

**Figure B.2.**
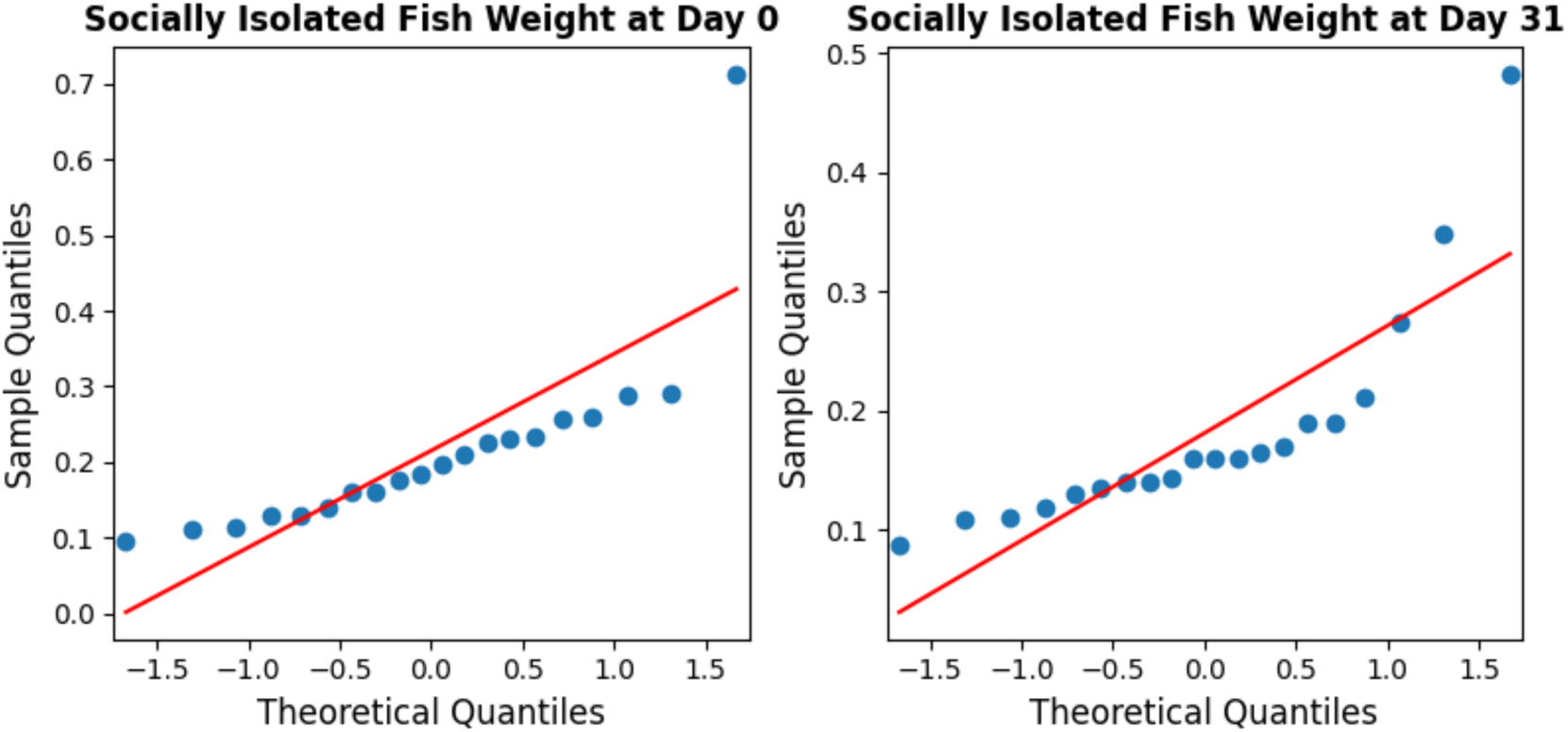
QQ plots for body weight day 0 (left) and day 31 (right) in socially isolated fish. These plots were used to visually assess the normality assumption for the change-score analysis.

### Control Fish

**Figure B.3.**
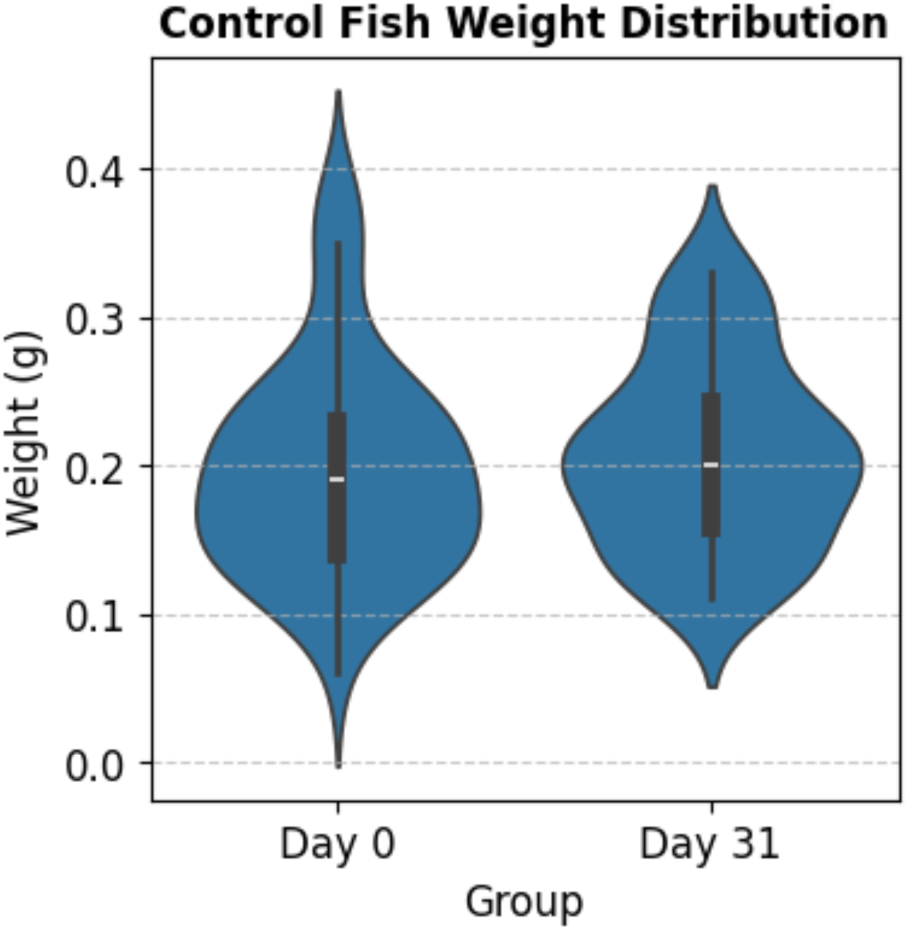
Violin plot for the distribution of body weight in day 0 and day 31 in the control group.

**Figure B.4.**
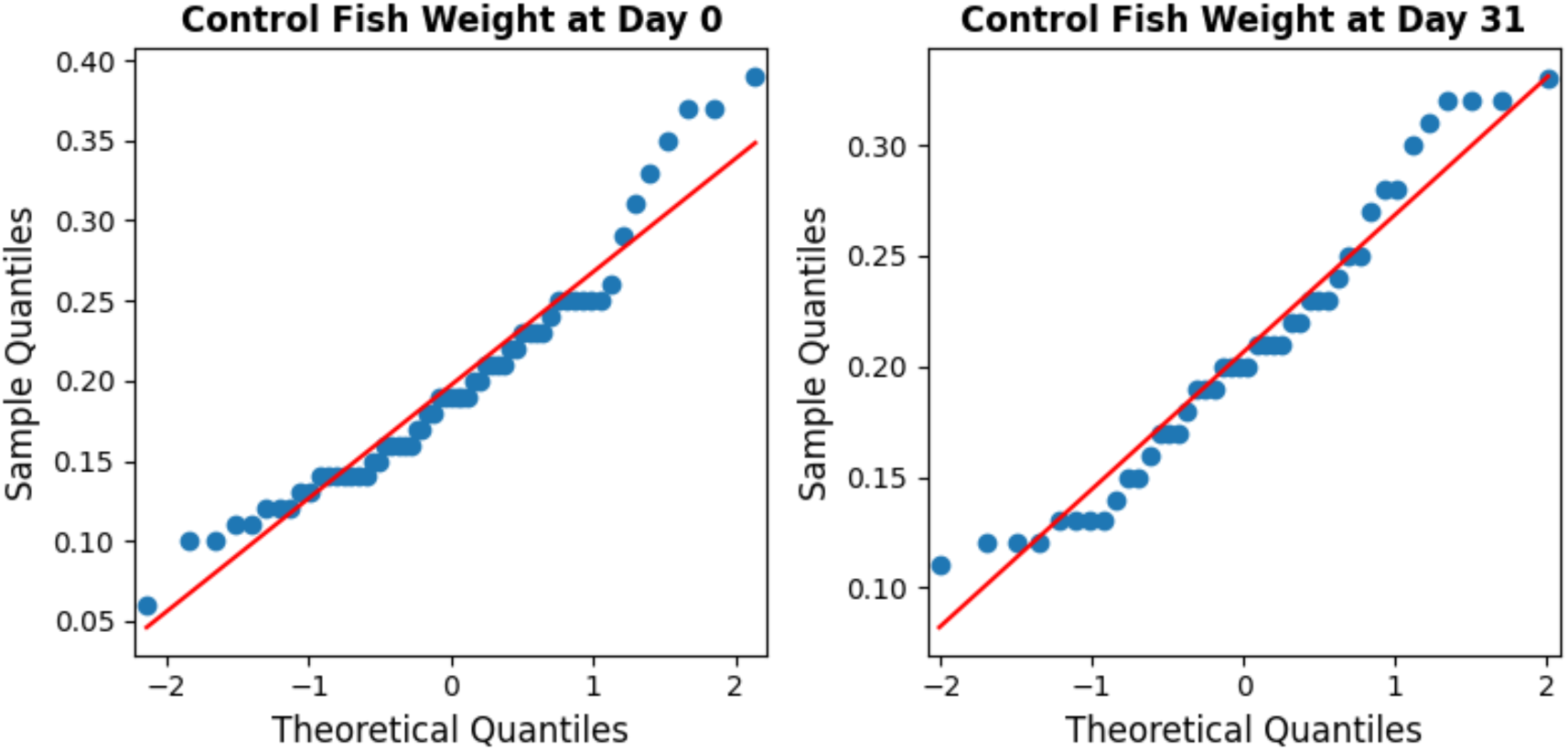
QQ plots for body weight day 0 (left) and day 31 (right) in socially isolated fish. These plots were used to visually assess the normality assumption for the change-score analysis.

## Appendix C: Social Preference and Movement Speed Measures

**Figure C.1.**
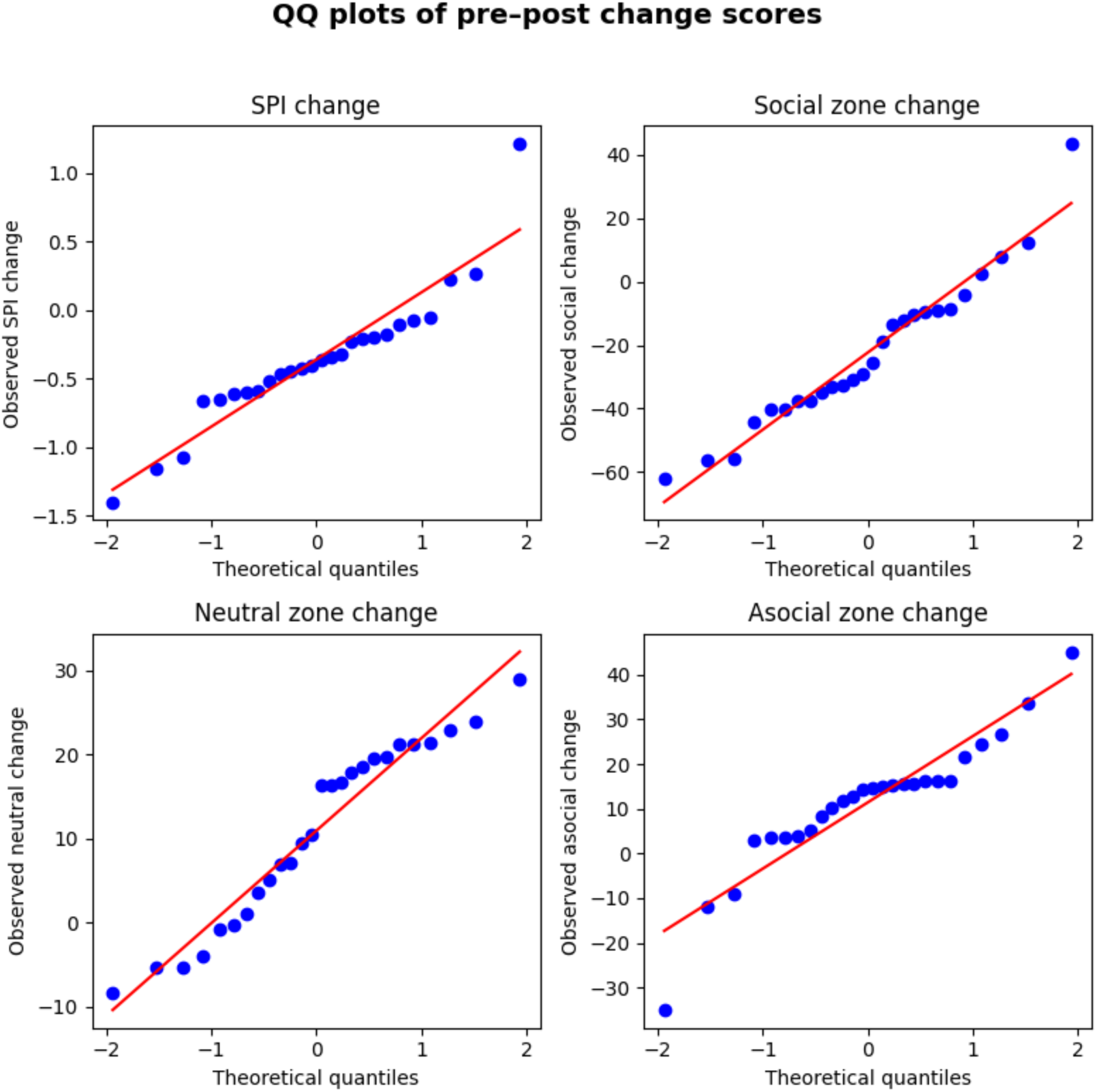
QQ plots for Day 0 to Day 31 changes in SPI (top-left), social-zone (top-right), neutral-zone (bottom-left), and asocial-zone (bottom-right). These plots were used to visually assess the normality assumption for the change-score analysis.

## Appendix D: Social Isolation Protocol

### Day-0

| Task<br>(check two fish<br>per day) | Method |
| --- | --- |
| Social Interaction | 1. Each test fish was individually introduced to the central zone of the testing apparatus and was temporarily separated by a transparent dividing door that could be slid up. This door divides the central zone from the left and right arms of the corridor.<br>2. 30 seconds of acclimatisation in the central part with the glass doors shut.<br>3. Following the initial 30-second acclimation interval, the two sliding dividers should be gently lifted, and the zebrafish should be released to explore the apparatus for 5 minutes.<br>4. Alternate the side on which the fish shoal was put and conduct the trial for the target fish again. |
| Dissolved oxygen<br>consumption rate | 1. Ensure the tank is filled with water up to its brim.<br>2. Put the Zebrafish in the inner tank.<br>3. Dip the respirometer in the inner tank.<br>4. Add parafilm to ensure that the enclosure is *airtight*<br>5. Check the reading on the respirometer every minute and note it down every five minutes for 90 minutes. |
| Feeding | ad libitum Tetra-Tetramin flakes once daily |

### Day-31

| Task | Method |
| --- | --- |
| Social Isolation | 1. Use opaque white containers, fill them with embryo water, and keep one fish per container.<br>2. 2 fish were socially isolated each day for 30 days |
| Social Interaction | 1. Each zebrafish was individually introduced to the testing apparatus's central zone, temporarily separated by a transparent dividing door that could be slid up. This door divided the central zone from the left and right arms of the corridor. |
|  | 2. 30 seconds of acclimatisation in the central part with the glass doors shut.<br>3. Following the initial 30-s acclimation interval, the two sliding dividers should be gently lifted, and the zebrafish should be released to explore the apparatus for 5 min.<br>4. Alternate the side on which the fish shoal was put and conduct the trial for the target fish again. |
| Dissolved oxygen consumption rate | 1. Ensure the tank is filled with water up to its brim.<br>2. Put the test fish in the inner tank.<br>3. Dip the respirometer in the inner tank.<br>4. Add parafilm to ensure that the enclosure is *airtight*<br>5. Check the reading on the respirometer every minute and note it down every five minutes for 90 minutes. |
| Feeding | ad libitum Tetra-Tetramin flakes once daily |

## Appendix E: Procedure to make embryo water

Social isolation was carried out in embryo water to remove olfactory cues present in the water. The embryo water was prepared using the following protocol for a 60x stock developed by the Cold Spring Harbour Laboratory (2011) for the use of zebrafish embryos:

a. Dissolve the following salts in 2L of H20: 34.8 g NaCl; 1.6 g KCl; 5.8 g CaCl2·2H2O; 9.78 g MgCl2·6H2O.
b. Adjust the pH to 7.2 with NaOH.
c. Autoclave.
d. To prepare 1X medium, dilute 16.5mL of the stock to 1 L
e. Add 100 µL of 1% methylene blue (Sigma-Aldrich)

The 1x solution was prepared freshly from the stock in small batches as needed.

